# Molecular mechanism regulating transcriptional control of the *hig* toxin-antitoxin locus of antibiotic-resistance plasmid Rts1 from *Proteus vulgaris*

**DOI:** 10.1101/2021.03.04.434028

**Authors:** Ian Pavelich, Marc A. Schureck, Dongxue Wang, Eric D. Hoffer, Michelle Boamah, Nina Onuoha, Stacey J. Miles, C. Denise Okafor, Christine M. Dunham

## Abstract

Regulation of ubiquitous bacterial type II toxin-antitoxin (TA) gene pairs occurs via a negative feedback loop whereby their expression is typically responsive to changing levels of toxins at the transcriptional level similar to a molecular rheostat. While this mechanism can explain how certain TA complexes are regulated, accumulating evidence suggests diversity in this regulation. One system for which the negative feedback loop is not well defined is the plasmid-encoded HigBHigA TA pair originally identified in a post-operative infection with antibiotic resistant *Proteus vulgaris*. In contrast to other type II TA modules, each *hig* operator functions independently and excess toxin does not contribute to increased transcription *in vivo*. Structures of two different oligomeric complexes of HigBHigA bound to its operator DNA reveal similar interactions are maintained suggesting plasticity in how *hig* is repressed. Consistent with this result, molecular dynamic simulations reveal both oligomeric states exhibit similar dynamics. Further, engineering a dedicated trimeric HigBHigA complex does not regulate transcriptional repression. We propose that HigBHigA functions via a simple on/off transcriptional switch regulated by antitoxin proteolysis rather than a molecular rheostat. The present studies thus expand the known diversity of how these abundant bacterial protein pairs are regulated.

**IMPORTANCE:** Bacteria respond to various stimuli by rapidly regulating gene expression to control growth. The diversity in how bacteria inhibit growth is exemplified by the abundance and diversity of toxin-antitoxin (TA) gene pairs. To tightly regulate their own expression, antitoxin proteins function as transcriptional autorepressors with additional regulation imparted by responsiveness of the system to toxin concentrations, similar to a molecular rheostat. However, some TAs do not appear to be responsive to changing levels of toxin. To expand our understanding of diverse TAs, we studied the regulation of a structurally distinct TA called <u>h</u>ost inhibition of <u>g</u>rowth (HigBA) originally discovered on the antibiotic resistance Rts1 plasmid associated with *Proteus vulgaris*. We find that the *hig* operon is regulated via a simple on/off transcriptional switch that is incalcitrant to changing toxin levels. These results expand the known mechanistic diversity of how TA pairs regulate their expression.

## INTRODUCTION

Bacterial toxin-antitoxin (TA) genes are bicistronic operons found in mobile genetic elements and bacterial chromosomes (1–3). Type II TA modules consist of toxin and antitoxin protein components that form architecturally diverse macromolecular complexes in the absence of external stimuli and during nutrient-rich growth. Although these gene pairs were first identified on plasmids and in bacteriophages (4–9), TAs are highly abundant in free-living bacteria where they appear to have different functions. In their role in plasmid maintenance, the toxin component can induce post-segregational killing if both genes are not inherited (6). In the past few years, conflicting experimental data on the endogenous activities of TAs have led to ambiguity and controversy surrounding their roles in bacterial physiology (10).

Regulation of type II TA pairs frequently occurs at the transcriptional level via a negative feedback loop (10–12). Antitoxin proteins contain a DNA-binding motif and repress at operator sites that overlap with promoters of TA genes. Toxin proteins are either recruited to their cognate antitoxins bound at these operator sites or bind to operators as TA complexes where they function as co-repressors, allowing the system to be responsive to changes in toxin expression levels. Further, TAs can form different oligomeric complexes when bound at operator sites that alter their binding thermodynamics to result in a gradient of the transcriptional response similar to a molecular rheostat (13, 14). This response gradient can also be influenced by cooperative TA binding at adjacent operator sites; this process is known as “conditional cooperativity” (15–18). However, this model is dependent on the architectural organization of TAs pairs which can diverge significantly (12, 19). Thus, it is unclear if conditional cooperativity can explain the regulation of all type II TA systems.

The structural diversity and distinct toxin- and DNA-binding motifs of different type II antitoxin proteins may partially explain why they can exert different mechanisms of autoregulation (11). Antitoxins contain ribbon-helix-helix (RHH), helix-turn-helix (HTH), Phd/YefM or SpoVT/AbrB DNA-binding motifs, with RHH and HTH being the most common (20–26). The type of DNA binding motif affects transcriptional repression. HTH-containing antitoxins contain a complete DNA-binding motif while RHH-containing antitoxins contain a half site requiring antitoxin dimerization for DNA binding. TA operons usually contain multiple operator sites and antitoxin binding at adjacent sites can lead to cooperativity and an increase in transcriptional repression (17, 18, 27). Antitoxins are particularly susceptible to proteases especially during changing cellular conditions (28). This reduction in antitoxin concentration increases free toxin levels that when free, can inhibit growth. Free toxin can also interact with antitoxins bound at their operators changing the oligomeric state of the TA complex during repression. These oligomeric state changes, in turn, can lead to differences in the ability of TAs to bind their operator and influence the extent of repression; effectively this responsiveness allows the system to function as a molecular rheostat (**Fig. 1**). In contrast, other TA systems do not appear to be responsive to changing levels of toxin and instead are simple on/off transcriptional switches (29, 30). While there exists some experimental evidence that distinguishes between the molecular rheostat and the on/off switch modes of regulation, at present the molecular basis for each mechanism is ambiguous because there is little or no structural evidence as a foundation for each model.

**Figure 1.**
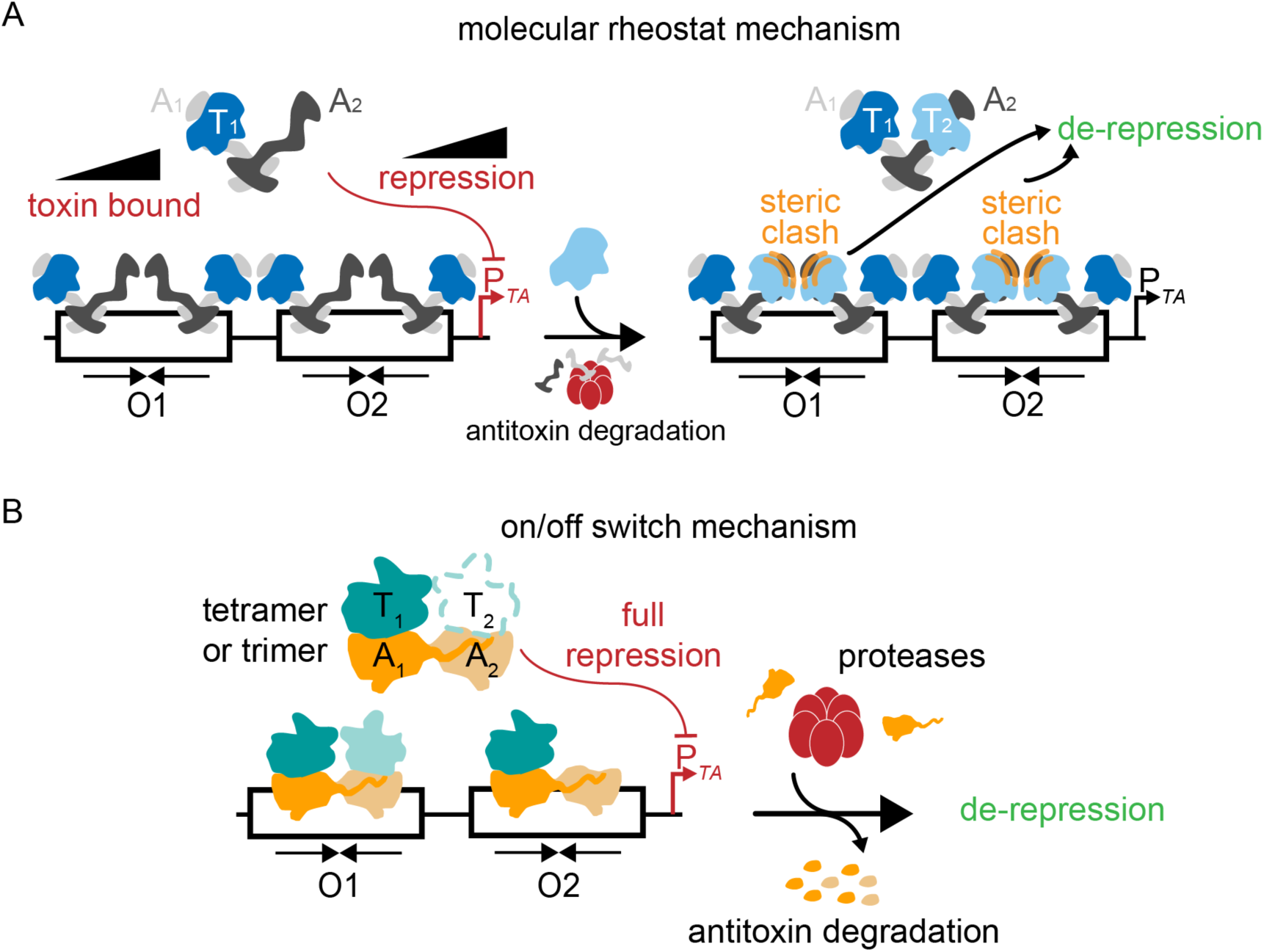
Diverse transcriptional control mechanisms that regulate expression of toxin-antitoxin complexes. Toxin (T) and antitoxin (A) proteins form multimeric complexes that bind operator sites (O1 and O2) that overlap with their promoters (P*_TA_*) to repress transcription. (**A**) In some type II toxin-antitoxin systems, changing levels of toxins (due to antitoxin proteolysis) that bind to the repressor complex leads to steric clashes and/or changes in affinity causing de-repression. In this case, the system functions as a molecular rheostat responsive to toxin levels. (**B**) In contrast, other toxin-antitoxin systems are not sensitive to changes in toxin concentrations and thus function as on/off transcriptional switches dependent on antitoxin depletion from proteolysis.

The *<u>h</u>ost inhibition of <u>g</u>rowth* BA (*higBA*) TA module seems to function as an on/off switch. The HigBHigA pair was first identified on the antibiotic-resistance plasmid Rts1 associated with *Proteus vulgaris* and discovered post-operatively in an urinary tract infection (31, 32) (we call this TA pair “HigBHigA” to denote both the HigB toxin and HigA antitoxin proteins). The HigB toxin belongs to the RelE family of toxins, resembles a microbial ribonuclease and cleaves mRNA substrates bound to a translating ribosome (26, 33–35). Although there are HigBHigA TA pairs found chromosomally and these HigB toxins are also RelE family members (36, 37), the structural organization and the regulation of these systems compared to the *P. vulgaris* associated module seems to be different (26). While all known HigA homologs contain a HTH DNA-binding motif, the *P. vulgaris* associated antitoxin binds to each of its operator sites (O1 and O2) in a non-cooperative manner (26, 38). Here, we test the two models of transcriptional regulation, rheostat versus on/off switch, to determine the molecular mechanism of action of HigBHigA.

## RESULTS

### Transcriptional repression at each operator is independently regulated

The *hig* promoter (P*hig*) is negatively autoregulated by the HigBHigA complex binding at operators O1 and O2 that overlap with the -35 and -10 promoter sites (32) (**Fig. 2A**). The HigBHigA complex forms a tetrameric assembly with two HigB monomers and a HigA dimer (HigB_2_HigA_2_) (26). Each HigA antitoxin contains a single HTH DNA-binding motif and forms an obligate dimer, meaning that two HigA antitoxins in one HigB_2_HigA_2_ complex bind two inverted repeats of a single DNA operator (26, 38). To determine if we could build oligomeric complexes *in vitro*, we monitored the binding of the HigB_2_HigA_2_ complex to *hig* (O1 and O2) using an electrophoretic mobility shift assay (EMSA) (**Fig. 2B**). The HigB_2_HigA_2_ complex was purified according to previously published protocols and the DNA probe used in the EMSA consists of the entire 61 basepair (bp) operator region (**Table S1**). Titration of HigB_2_HigA_2_ with a constant amount of *hig* causes two molecular weight shifts, indicating binding of HigB_2_HigA_2_ at each operator site (**Fig. 2B**, top). To determine whether HigB_2_HigA_2_ binds with a higher affinity to either O1 or O2, all 21 nucleotides in each operator were randomized individually (38). Each of these 21 nucleotides located in either O1 or O2 were previously shown to be protected upon HigA binding (39). Therefore any change in the mobility of the DNA band using a scrambled O1 or O2 would represent binding of HigB_2_HigA_2_ to a single operator. HigB_2_HigA_2_ binds to each of the two sites represented as a single molecular weight shift and both result in similar dissociation binding constants (0.36 ± 0.09 μM for O1 and 0.24 ± 0.04 μM for O2) (**Fig. 2B**, middle and lower; **Table S2**). These data indicate that HigB_2_HigA_2_ recognizes each operator independently to form a high affinity interaction. This observation appears to be an important distinction from other type II TAs where TA complexes binding at an operator influences the binding of TAs at adjacent operators (**Fig. 1**).

**Figure 2.**
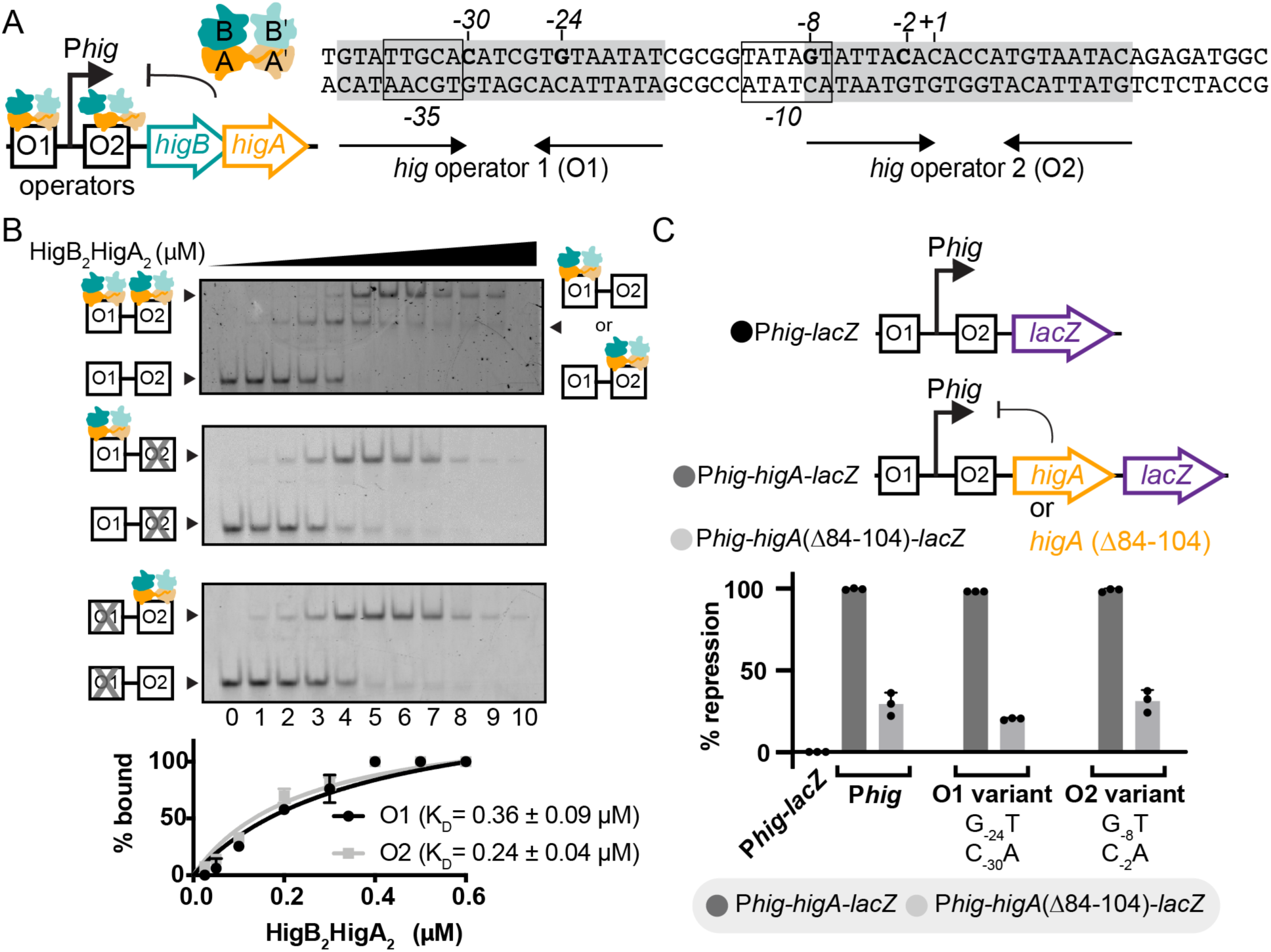
Binding of HigB_2_HigA_2_ to a single operator is sufficient for transcriptional repression of the *hig* operon. (**A**) *Left*, organization of the *hig* operon containing the operators O1 and O2, the P*hig* promoter, *higB* toxin and *higA* antitoxin genes. *Right*, the nucleotide sequences of O1 and O2, with the +1 transcriptional start site and the -35 and - 10 sites indicated. The sequence recognized by HigA is shown in grey and operator nucleotides C_-30_, G_-24_, G_-8_, and C_-2_ important for HigA binding are shown in bold. (**B**) EMSA of HigB_2_HigA_2_ binding to wild-type P*hig* (top), O1 only (O2 scrambled; middle), and O2 only (O1 scrambled; bottom) DNA. Band intensities were plotted from EMSAs as the percent of HigB_2_HigA_2_ bound to DNA versus HigB_2_HigA_2_ (concentrations used: 0, 0.025, 0.05, 0.1, 0.2, 0.3, 0.4, 0.5, 0.6, 0.7, 0.8 μM). Curves represent the fit from which K_D_s were calculated. (**C**) β-gal assays of *E. coli* BW25113 transformed with pQF50-P*hig*-*lacZ* (black), pQF50-P*hig*-*higA*-*lacZ* (dark grey), or pQF50-P*hig*-*higA*(Δ84-104)-*lacZ* (light grey). Each operator site was tested using known operator mutations of either O1 (G_-24_T, C_-30_A) or O2 (G_-7_T, C_-2_A) (38).

To test whether HigB_2_HigA_2_ binding at a single operator results in transcriptional repression *in vivo*, we designed a series of constructs that encode *lacZ* in three different contexts: downstream of the P*hig* promoter (pQF50-P*hig*-*lacZ*), downstream of wild-type HigA (pQF50-P*hig*-*higA*-*lacZ*), or downstream of a C-terminally truncated HigA (pQF50- P*hig*-*higA*(Δ84-104)-*lacZ*) (**Fig. 2C**). The pQF50-P*hig*-*lacZ* construct reports on the activity of P*hig* in the absence of repressor HigA, while the HigA-encoding pQF50-P*hig*-*higA*-*lacZ* construct reports on how expression of HigA represses P*hig*. The third construct serves as a control as the HigA(Δ84-104) variant is unable to dimerize and bind P*hig* but is comparably expressed (26, 38). P*hig*-*lacZ* shows robust β-galactosidase (β-gal) activity which we normalize to represent complete transcriptional repression (this is represented as 0% repression; **Fig. 2C**). Expression of HigA efficiently represses P*hig*, whereas HigA(Δ84-104) restores P*hig* activity to near P*hig*-only levels, indicating transcriptional repression is dependent on HigA expression and dimerization (**Fig. 2C**). To test whether HigA_2_ binding to a single operator region is sufficient for repression, we mutated either O1 or O2 recognition sequences important for HigA binding (38) and repeated the previously described β-gal assays. Mutations to either O1 (G_-24_T/C_-30_A) or O2 (G_-8_T/C_-2_A) do not affect HigA_2_ repression at a single mutant P*hig* when compared to the wild-type P*hig* (**Fig. 2C**, compare middle bars in “O1 variant” and “O2 variant” group to “P*hig*”). These results show that HigA_2_ binding at either O1 or O2 is sufficient for repression (data also presented in Miller Units in **Fig. S1**). Further, these data indicate that HigA_2_ binding at each *hig* operator is not cooperative, and that a single operator is sufficient for transcriptional repression.

### Addition of HigB destabilizes interactions between HigA and DNA *in vitro*

The expression of most type II TA complexes is regulated at the transcriptional level by changing toxin and antitoxin concentrations as a result of increased proteolysis of the antitoxin during external stimuli (17, 18, 22). We reasoned that the HigBHigA complex may transcriptionally repress using a different mechanism because of the structural diversity of the HigA antitoxin. The unique structure of HigA could influence its interactions with *hig* or HigB toxin binding. For example, the HigA HTH motif is a complete DNA-binding domain and by extension, the obligate HigA_2_ dimer contains two DNA-binding domains. Other type II antitoxins typically contain a single DNA-binding motif formed by antitoxin dimerization (**Fig. 1**). These differences are likely important for the changing of different oligomeric TA complexes bound to operators allowing the system to respond to changing toxin levels. To test this, we explored whether the presence of excess HigB changes the molecular interactions of HigA with the O1-O2 operators (**Fig. 3**). Addition of HigA_2_ to O1-O2 results in two shifted bands of lower mobility, indicative of complexes with a single HigA_2_ dimer bound to one operator (**Fig. 3A**, **yellow**), or two HigA_2_ dimers binding to each operator (**Fig. 3A, orange**). Increasing amounts of HigB results in the formation of two additional higher molecular weight species (**Fig. 3A**). In this case, either the HigB monomer binds to each of the HigA_2_ dimers to form a trimeric HigBHigA_2_ complex (**Fig. 3A, red**) or both HigB monomers bind to a single HigA_2_ dimer as a tetrameric HigB_2_HigA_2_ (**Fig. 3A, purple**). As the HigA_2_ dimer is already bound at both O1 and O2, it is unlikely that the molecular weight species would represent occupation of only a single operator site. While the third shift may contain a mixed population of oligomeric states where HigB binds only one HigA_2_ dimer (**Fig. 3A, red**), an additional observable migration shift may be difficult to observe. Thus, we assume the slowest migrating band is a tetrameric HigB_2_HigA_2_-O1-O2 complex. When the molar ratio of HigB to HigA exceeds one (two HigB monomers to one HigA_2_ dimer or 1.0 μM HigB for 0.5 μM HigA_2_ dimer), only free DNA is observed suggesting that excess HigB destabilizes the association between HigA and DNA. These results appear to be similar to what has been observed with other type II TA complexes that exhibit regulation via conditional cooperativity (17, 18, 27).

**Figure 3.**
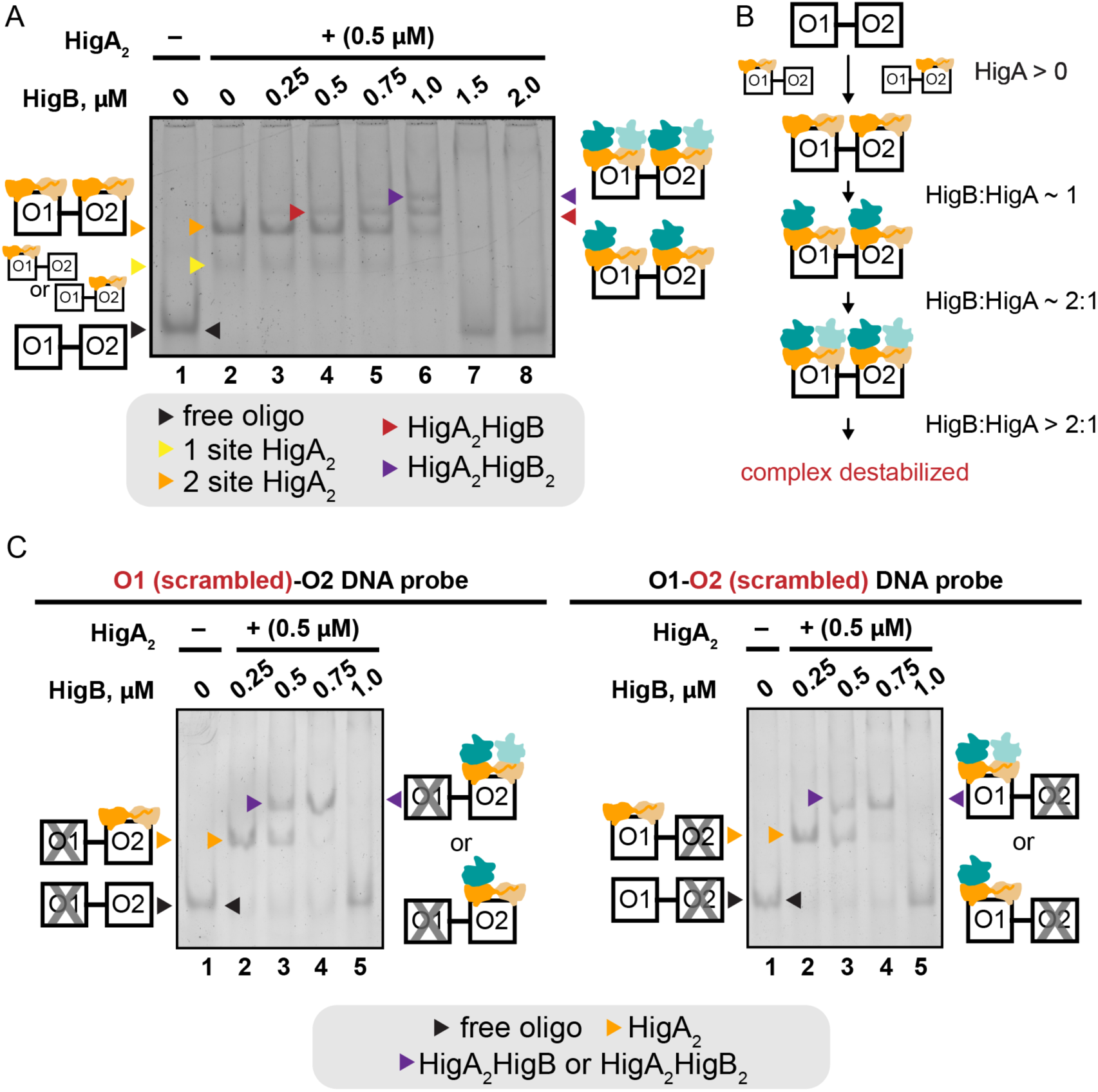
Formation of higher oligomeric states upon addition of HigB to the HigA-DNA complex. (**A**) EMSA of HigB addition to the HigA_2_-O1-O2 DNA complex. The O1-O2 DNA migrates the fastest (lane 1, black arrowhead) and incubation with the 0.5 µM HigA_2_ dimer results in a retardation of O1-O2 to form two shifts (lane 2, yellow and orange arrowheads). Increasing amounts of HigB results in the formation of higher molecular weight complexes (lanes 3-6, red and purple arrowheads). When the molar ratio of HigB to HigA exceeds 1 (>1.0 μM HigB for 0.5 μM HigA_2_ dimer), the high molecular weight shifts are no longer observed indicating that neither HigA nor HigB binds. (**B**) Schematic of the different HigBHigA-O1-O2 DNA complexes formed with changing the HigB:HigA ratio. Multiple oligomeric arrangements are possible in the case of a trimeric HigA_2_HigB complex but only one example is shown. (**C**) EMSA of HigB addition to the HigA_2_-O1-O2 DNA complex when either O1 or O2 is scrambled. O1-O2 DNA migrates the fastest (lane 1, black arrowhead) and incubation with the HigA_2_ dimer results in a retardation of O1-O2 to form two shifts (lane 2, orange arrowhead). The addition of HigB results in the formation of higher molecular weight complexes (lane 3, purple arrowhead) with a molar excess of HigB over HigA causes the scrambled DNA probe to be released (lane 5).

To determine whether the instability of the HigBHigA complex with O1-O2 is dependent upon the occupancy of both O1 and O2, we next tested complex formation on DNA containing either a scrambled O1 or O2 to prevent HigA_2_ binding (same mutation as used in Fig. 2). Addition of HigA_2_ to DNA containing either a scrambled O1 or O2 results in a slower moving species indicative of the HigA_2_ dimer binding at one operator (**Fig. 3C, orange**). Increasing amounts of HigB shows a single, slower moving molecular weight species (**Fig. 3C, purple**). At this point, we assume the HigBHigA complex is tetrameric (HigB_2_HigA_2_) given the prior crystal structures (26). Once the HigB to HigA molar ratio almost exceeds 1 (i.e. 1.0 μM HigB for 0.5 μM HigA_2_ dimer), the complex is unstable and DNA is released. Interestingly, in the context of both O1 and O2 (**Fig. 3A**), the molar ratio of HigB to HigA needs to exceed 1 however, when only one operator is available, this release of DNA occurs at slightly less then 1 molar equivalents of HigB and HigA. Therefore, the instability of HigBHigA binding a single operator appears to be accelerated in the absence of an adjacent HigBHigA complex.

### Structure of HigB_2_HigA_2_-O2 DNA

To determine how HigBHigA interacts with its operator DNA, we pursued a high-resolution X-ray crystal structure of HigBHigA bound to a single operator, O2. We performed crystallization trials using two HigBHigA constructs: a six histidine (His_6_) affinity tag located at the N terminus of HigB and a His_6_ affinity tag located at the C terminus of HigA. Both HigBHigA variants crystallized in the same condition, however, each resulted in a different oligomeric state of the HigBHigA complex bound to O2. The HigBHigA-His_6_-O2 complex crystalized in the monoclinic space group C121, was determined to 2.4 Å resolution by single wavelength anomalous diffraction phasing and contained a HigA_2_ dimer bound to two HigB monomers (**Fig. 4**). The His_6_-HigBHigA-O2 complex crystalized in the tetragonal space group I4_1_, was determined by molecular replacement using the previously determined HigA_2_ model (PDB code 6CF1) to 2.8 Å resolution and contained a HigA_2_ dimer bound to a single HigB (**Fig. 5**; **Table S3**). In both structures, residues 1–91 were built for each HigB monomer (92 total residues) and all nucleotides (1–21) were built for the O2 DNA duplex (**Fig. S2**). Residues 1–101 and 1–102 in the HigB_2_HigA_2_ structure and residues 1–91 and 1–95 in the HigBHigA_2_ structure were modeled (104 total residues) (**Fig. S2**).

**Figure 4.**
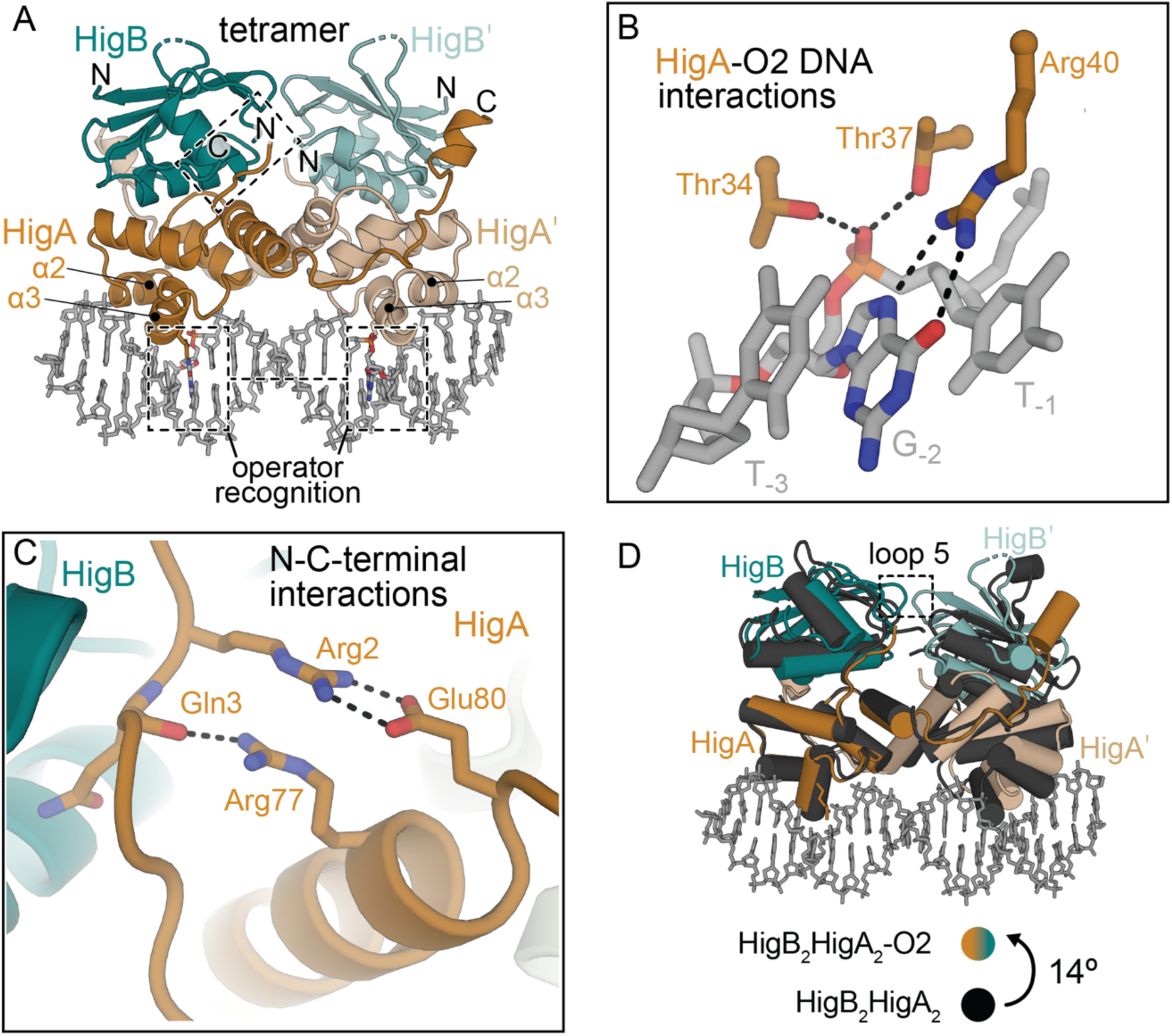
Structure of tetrameric HigB_2_HigA_2_ bound to O2 DNA. (**A**) The 2.4-Å structure of tetrameric HigB_2_HigA_2_-O2 DNA complex (PDB code 6W6U). HigA recognizes the T_-1,_ G_-2,_ T_-3,_ A_-4_ DNA operator region via α2 and α3. N and C-terminal regions of HigA are boxed. (**B**) HigA Arg40 makes the only sequence specific interactions with the nucleobase of G_-2_ while HigA residues Thr34 and Thr37 (both from α3) stably interact with the phosphate of G_-2_. (**C**) The N- and C-terminal residues of HigA become ordered upon both HigB binding. HigA residue Arg77 forms a hydrogen bond with the backbone carbonyl of Gln3 and Arg2 and Glu80 interact via a salt bridge. (**D**) Comparison of the tetrameric HigB_2_HigA_2_ complex (all black; PDB code 4MCX) and HigB_2_HigA_2_-O2 DNA complex (PDB code 4MCX) reveal a ∼14° rotation of HigB_2_HigA_2_ away from DNA that allows recognition.

**Figure 5.**
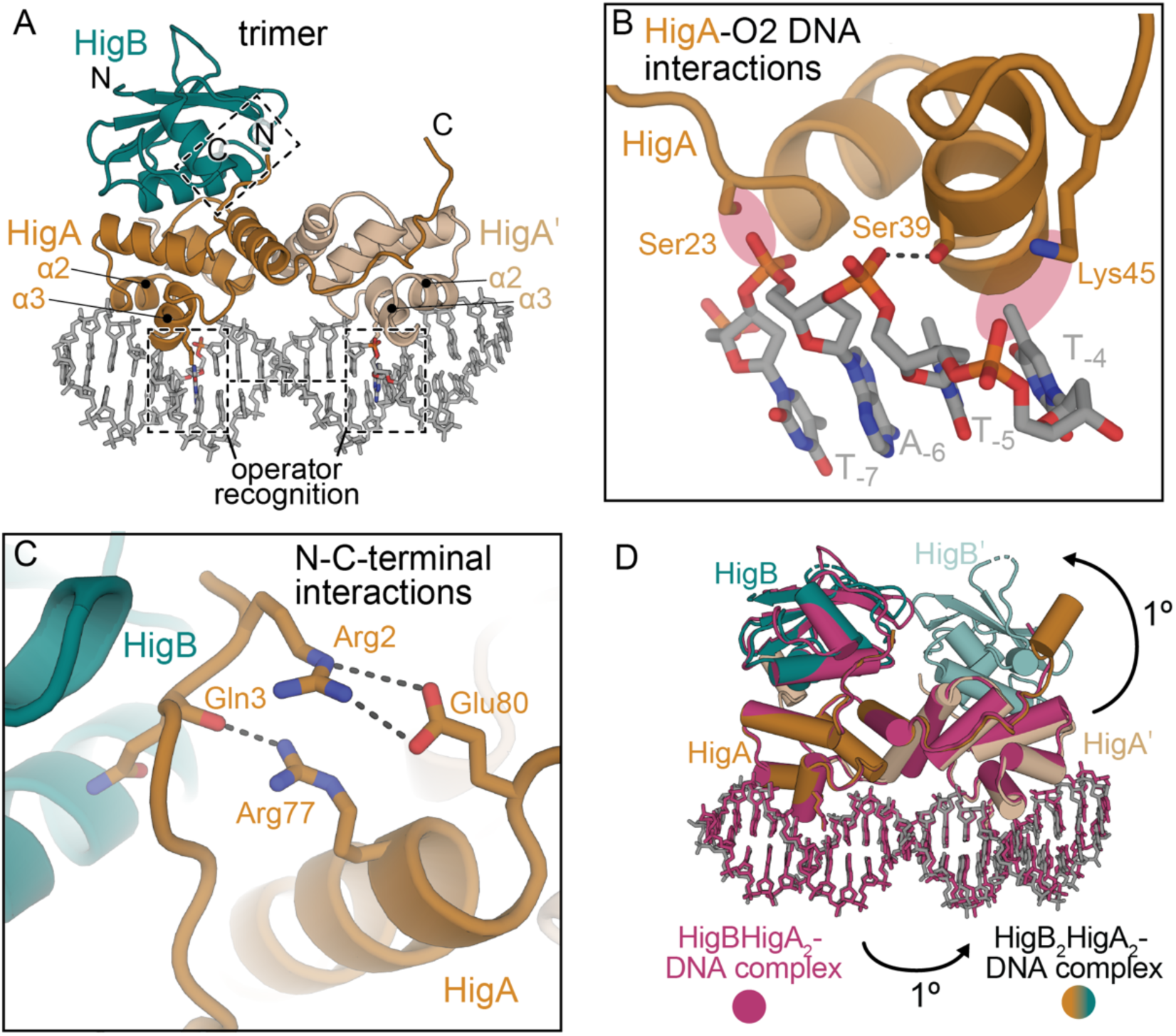
Structure of trimeric HigBHigA_2_ bound to O2 DNA. The 2.8-Å structure of trimeric HigBHigA_2_-O2 DNA (PDB code 6WFP). HigA recognizes the T_+6,_ G_+7,_ T_+8,_ A_+9_ DNA region via α2 and α3. N and C-terminal regions of HigA are boxed. (**B**) In the HigB_2_HigA_2_- O2 DNA structure (PDB code 6W6U), HigA residues Ser23, Ser39, and Lys45 interact with the backbone phosphate of T_-7_, T_-5_, and T_-4_ respectively to rigidify the T_-1,_ G_-2,_ T_-3,_ A_-4_ sequence for nucleotide-specific recognition on the opposite strand. In the trimeric HigBHigA_2_-O2 structure, only Ser39 interacts with the phosphate backbone and Ser23 and Lys45 are too distant (red highlighted region). (**C**) The N- and C-terminal residues of HigA become ordered upon a single HigB monomer binding similar to when two HigB monomers bind (Fig. 4C). (**D**) Comparison of trimeric HigBHigA_2_-O2 DNA (pink; PDB code 6WFP) and tetrameric HigB_2_HigA_2_-O2 DNA (PDB code 6W6U) are incredibly similar with an r.m.s.d of 0.7 Å (for 1479 equivalent atoms) and less than a ∼1° rotation.

The HTH motif in HigA consists of α2, loop 3 and α3 and this region interacts with the major groove of the operator O2 DNA (**Fig. 4A**). In the tetrameric HigB_2_HigA_2_-O2 structure, HigA contacts the T_-1,_ G_-2,_ T_-3,_ A_-4_ O2 sequence on the *hig* negative strand (38) (**Figs. 4A, B**). HigA residue Arg40 interacts with the Hoogsteen face of G_-2_ to make the only sequence-specific protein-DNA contact. Residues Thr34 and Thr37 (from α3) contact the phosphate of G_+7_ while the sidechains of Ser23 (from loop 2), Ser39 (from α3), and Lys45 (from α3) are all within hydrogen bonding distance of nucleotides T_-7_, A_-6_, T_-5_, and T_-4_, respectively which are located on the opposite DNA strand (**Fig. S3**). Additionally, Ala36 and Thr34 form van der Waals interactions with the nucleobase C5 methyl of A_-3_. These interactions are similar to those previously observed in the HigA_2_-O2 DNA interaction (38) and are also present between HigA and O2 on the opposite strand, indicating that HigB binding to form the tetrameric HigB_2_HigA2-O_2_ complex does not change interactions of HigA_2_ with O2.

The termini of antitoxins are typically intrinsically disordered contributing to their proteolysis during external stimuli. In the free HigA_2_ structure (38), the N terminus is disordered (**Fig. S4A,B**). Upon HigB binding, the HigA termini becomes ordered both in the free HigB_2_HigA_2_ structure (26) and upon binding DNA (HigB_2_HigA_2_-O2 DNA) (**Fig. 4C; Fig. S4C,D**). The N- and C-termini of HigA form intramolecular interactions in addition to interactions with α1 of an adjacent HigB in the crystal lattice (**Fig. S4A,B**). Specifically, N-terminal residues Arg2 (side chain) and Gln3 (backbone carbonyl) form salt bridges with C-terminal residues Glu80 and Arg77, respectively, and these interactions presumably stabilize the termini. Thus, binding of HigB stabilizes HigA both in the presence or absence of DNA.

Comparison of the overall architecture of HigB_2_HigA_2_-O2 DNA to HigB_2_HigA_2_ (26) or HigA_2_ (38) reveals subtle changes that may be important for O2 DNA binding and transcriptional repression. Aligning analogous HigA monomers from the HigB_2_HigA_2_-O2 and the HigB_2_HigA_2_ structures (PDB code 4MCX) reveals a ∼14° displacement of the adjacent, second HigA protomer (**Fig. 4D**). Similarly, comparison of the free HigA_2_ dimer (PDB code 6CF1) to HigB_2_HigA_2_-O2 also shows rotation of HigA upon DNA binding, although the movement is not as large as compared to when HigB is present (∼8° rotation versus a ∼14° rotation) (**Fig. S5**). Thus, HigA_2_ reorients to bind DNA and HigB binding to a HigA_2_-DNA complex minimally influences the protein-DNA interface.

### Structure of HigBHigA_2_-O2 DNA

As noted above, both the tetrameric HigB_2_HigA_2_-O2 DNA and trimeric HigBHigA_2_-O2 DNA crystal forms formed in the same crystallization conditions and resulted in two different macromolecular structures (**Table S3**). Interestingly, not all of the interactions seen in tetrameric HigB_2_HigA_2_-DNA are conserved in the trimeric HigBHigA_2_- O2 structure. While critical interactions of HigA with the T_-1,_ G_-2,_ T_-3,_ A_-4_ recognition sequence are maintained, α2 and α3 of the HTH DNA-binding motif slightly moves away from O2, no longer positioning Ser23 and Lys45 to hydrogen bond with the phosphates of T_-7_ and T_-4_ (**Fig. 5B; Fig. S3**).

Global comparison of the HigB_2_HigA_2_-O2 structure with the HigBHigA_2_-O2 structure reveal only a ∼1° difference emphasizing how similar the two structures are (**Fig. 5D**). Likewise there are very little differences in the position of HigA_2_ bound to O2 (38) in the absence or presence of HigB. Thus, it does not appear that HigB binding influences the position of HigA_2_ on DNA. It appears the largest structural change results from either HigA_2_ or HigB_n_HigA_2_ binding to DNA (∼14° rotation, where “n” denotes either a single HigB or two HigB monomers; **Fig. 4D**). Previously we described how HigA N-terminal residues Arg2 and Gln3 interact with its C-terminal residues Arg77 and Glu80 in the HigB_2_HigA_2_-O2 structure (**Fig. 4C**). We find that even a single HigB binding can cause these termini residues to become ordered (**Fig. S4E**).

A curious crystallization note for the trimeric HigBHigA_2_-O2 complex is that there is an adjacent molecule in the neighboring asymmetric unit that overlaps with the missing HigB (**Fig. S6**). This ejection of HigB from the HigBHigA complex is surprising given the known tight interactions of TA complexes where affinities are typically sub-nanomolar (29, 40–42). Therefore, we think it is unlikely that the trimeric HigBHigA_2_-O2 complex results from crystal packing. Interestingly, the structures of both HigBHigA complexes with the different placement of the His_6_ tag were solved and both found to be tetrameric HigB_2_HigA_2_ in the absence of DNA. Taken together, we propose that there is a mixture of both trimeric and tetrameric HigBHigA complexes bound to DNA in solution. We next sought to examine the functional relevance of the trimeric HigBHigA_2_-O2 complex.

### HigB_2_A_2_-O2 and HigBHigA_2_-O2 complexes exhibit similar dynamics

The structure of the trimeric HigBHigA_2_-O2 complex is intriguing as most models that describe the transcriptional regulation of type II TA systems conclude that such an oligomeric state is more stable than the fully loaded complex (18, 21, 42). However, prior to our new structure, there has been no biochemical or direct evidence for the existence of this oligomeric state. To assess the dynamics of both complexes in the presence or absence of O2, we performed molecular dynamics (MD) simulations of four complexes: HigB_2_HigA_2_ and HigBHigA_2_ in the presence or absence of O2 (**Fig. 6**). The trimeric HigBHigA_2_ complex in the absence of DNA has not been solved and we generated the model based upon the HigBHigA_2_-O2 structure. We obtained 1 microsecond-long MD trajectories of each complex and subsequently performed root mean square fluctuation (RMSF) analysis. This analysis reveals overall comparable dynamics: in tetrameric HigB_2_HigA_2_, binding to O2 only marginally affects dynamics, with the largest effects observed at intrinsically flexible regions such as the C termini of HigA monomers (residues 94-102) and loop 3 of HigB (residues 56-62) (**Fig. 6A**). In trimeric HigBHigA_2_, similar trends are observed, confirming that both oligomeric states represent similarly stable, DNA-bound complexes (**Fig. 6B**). One noted difference is that in the trimeric HigBHigA_2_-O2 complex, the C-termini of one of the two HigAs is disordered and is not modeled. Two HigB monomers binding causes the C-termini of HigA to regain order but while the C-termini of both HigA monomers can be modeled, this region still exhibits dynamic behavior.

**Figure 6.**
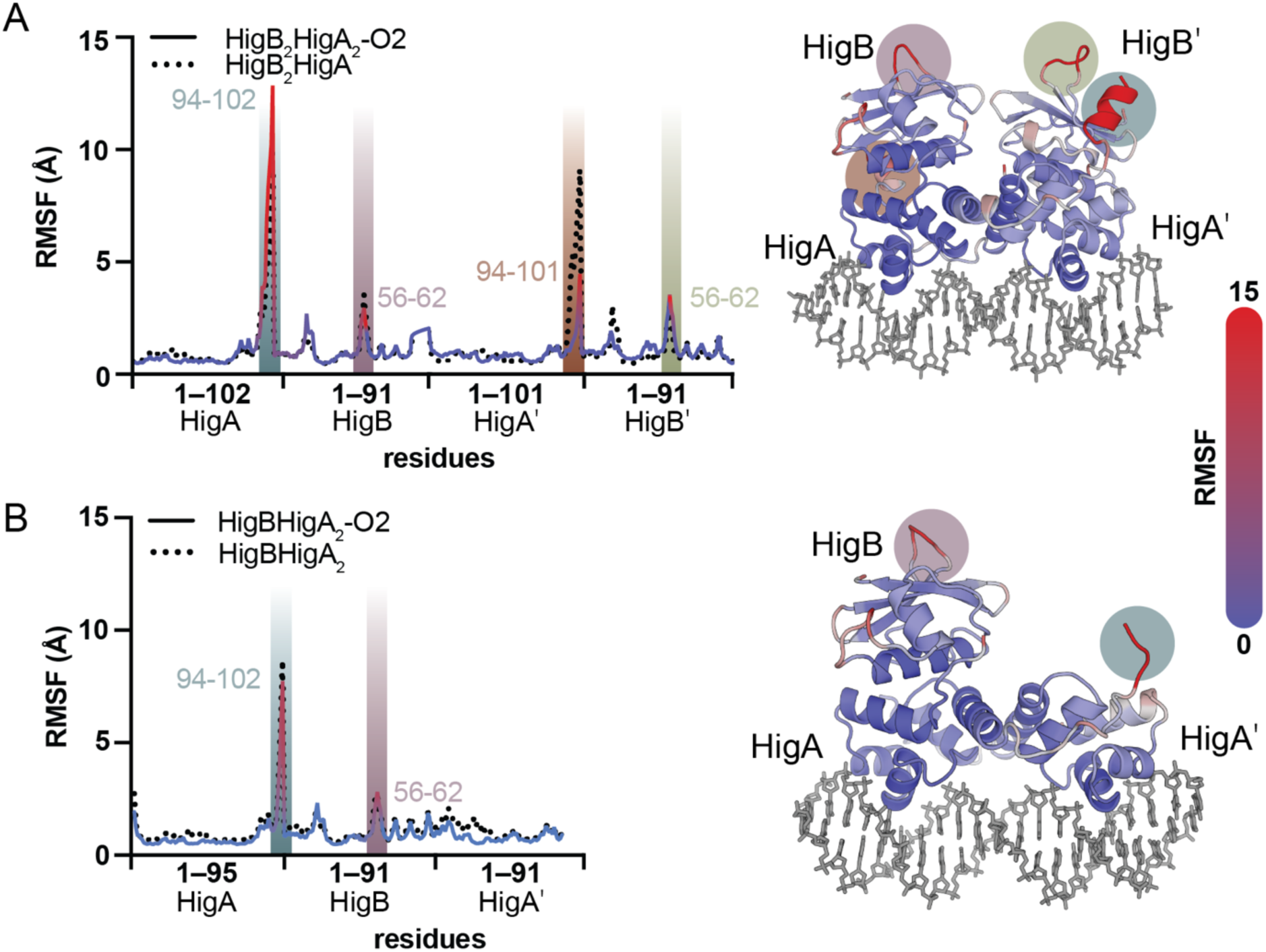
Trimeric HigBHigA_2_ and tetrameric HigB_2_HigA_2_ exhibit similar dynamics in the presence or absence of O2 DNA. Root-mean-square-fluctuations (RMSFs) of Cα atoms for each residue in the (**A**) HigBHigA_2_ or (**B**) HigB_2_HigA_2_ complexes are calculated from 1 ms MD trajectories. Regions that have increased RMSFs are indicated with highlighted bars that correspond to their positions on the HigBHigA-O2 structures (*right*). High RMSF spikes correlate to either labile C-termini of HigA or HigB loop regions with colored circles corresponding to the highlighted bars on the left.

### Engineered trimeric HigBHigA_2_ transcriptionally represses P*hig*

To test whether a trimeric HigBHigA_2_ complex represses transcription to the same extent as HigB_2_HigA_2_, we attempted to engineer such a variant. Comparison of the HigB_2_HigA_2_ structure with the HigB_2_HigA_2_-O2 DNA structure shows that the two HigB monomers move closer to each other to accommodate binding to DNA (**Fig. 7A**). In particular, HigB loop 5 (L5) located at the interface of the HigB monomers moves ∼4Å (**Fig. 4D**). We therefore extended L5 by the addition of a short, flexible sequence of four residues (Asn, Gly, Asn, Gly (NGNG); called HigB(L5ext)HigA_2_) to prevent concurrent binding of two HigB monomers to HigA_2_ (**Fig. 7A**). Expression and purification of HigB(L5ext)HigA_2_ showed a delayed elution of the complex from the size exclusion column as compared to wild-type HigB_2_HigA_2_ (**Fig. 7B**), at a volume corresponding to a molecular weight of 42 kDa (compared to 56 kDa for wild-type HigB_2_HigA_2_). The difference in apparent molecular weights indicates that the HigB(L5ext)HigA_2_ complex is ∼14 kDa smaller than the wild-type complex which roughly corresponds to a HigB monomer (molecular weight of ∼13 kDa). To assess its thermal stability, we performed nano-differential scanning fluorimetry (nano-DSF) which provides information on the melting temperature (T_m_) of the complex. Since this measurement is not at equilibrium, the inflection point is known as T_i_. HigB(L5ext)HigA_2_ is ∼5°C less thermostable than wild-type HigB_2_HigA_2_ (60.5°C vs. 54.0°C) consistent with an altered oligomeric state (**Fig. 7C**).

**Figure 7.**
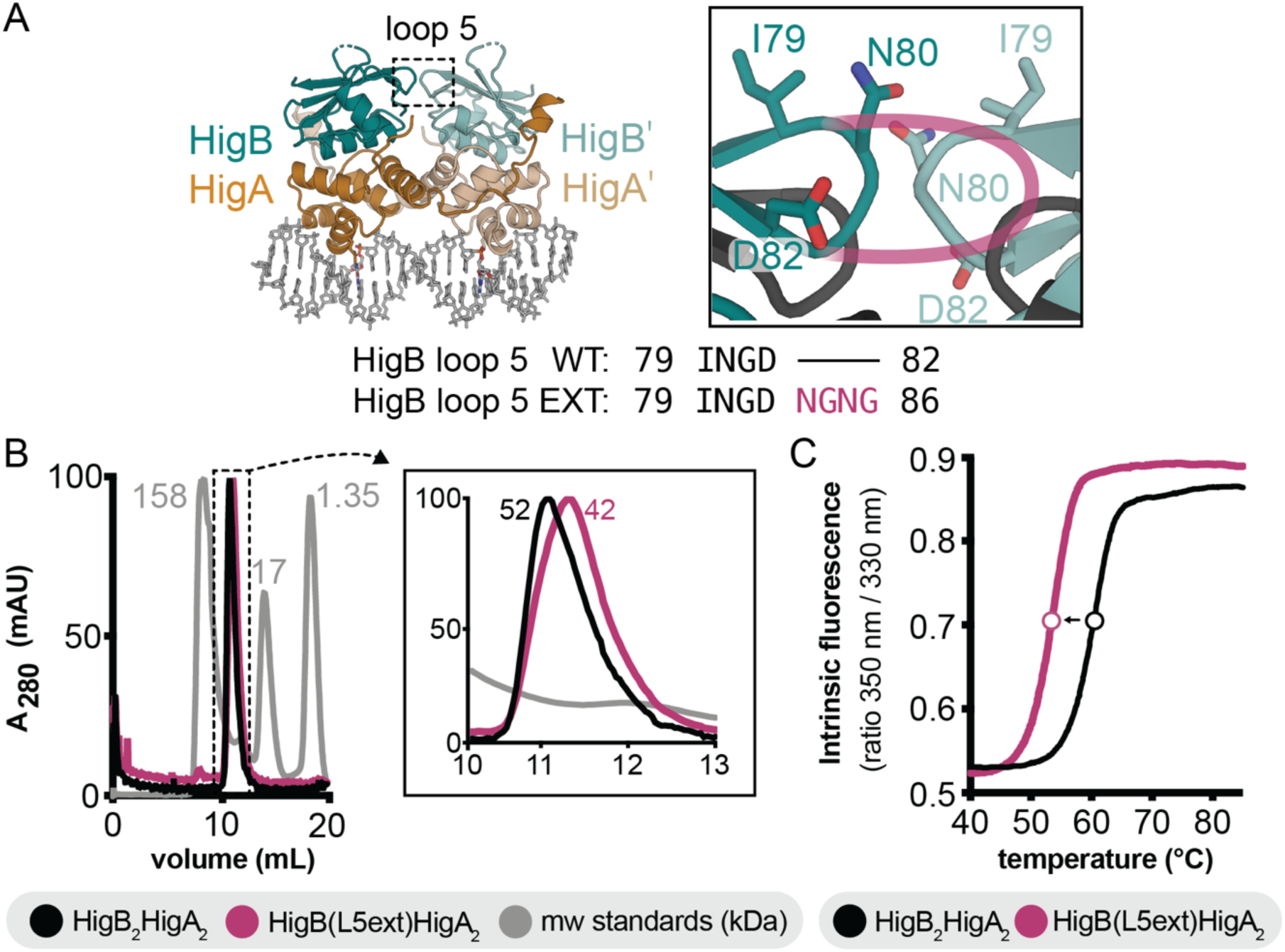
Engineering a trimeric HigBHigA_2_ complex. (**A**) To prevent two HigB monomers from binding, loop 5 (L5) of HigB was extended by the insertion of four residues (Asn, Gly, Asn, Gly; NGNG; “L5ext”, magenta) after residue Asp82. The dotted box indicating the L5 region of two HigB monomers is zoomed in (*right*). The theoretical extension of L5 is shown in magenta with the wild-type HigB and HigB(L5ext) amino acid alignment shown underneath. (**B**) Size exclusion chromatography of purified wild-type HigB_2_HigA_2_ shows an elution volume that corresponds to a molecular weight of 52 kDa. HigB(L5ext)HigA_2_ complex (magenta) elutes at a volume corresponding to a molecular weight of 42 kDa with the inset showing a zoomed in view. Molecular weight standards are shown in grey. (**C**) Nano-DSF analysis of wild-type HigB_2_HigA_2_ (black) and HigB(L5ext)HigA_2_ (magenta) shows that the HigB(L5ext)HigA_2_ complex has ∼5°C lower T_i_ value than HigB_2_HigA_2_. Fluorescence values were normalized to the highest tested temperature and the boundary of each line represents the mean ± SD of values of three independent experiments.

To test the ability of the HigB(L5ext)HigA_2_ variant to repress transcription at *hig*, we performed EMSA and β-gal assays using similar approaches as described above. The HigB(L5ext)HigA_2_ complex binds to both O1 and O2 DNAs at concentrations similar to wild-type HigB_2_HigA_2_ (**Fig. 8A**). Although an intermediate shift is observed (likely representing a HigA_2_ dimer bound), these data show the trimeric HigB(L5ext)HigA_2_ interacts with O1 and O2 in a similar manner as wild-type HigB_2_HigA_2_. To determine whether the HigB(L5ext)HigA_2_ complex represses P*hig in vivo*, we used the previously described P*hig* constructs and included a P*hig* containing the HigB(L5ext)HigA_2_ variant (pQF50-P*hig*-*higB*(L5ext)*higA*-*lacZ*; **Fig. 8B; Fig. S7**). As expected, P*hig* alone shows high β-gal activity (normalize to 0% repression) because of the absence of transcriptional repressor HigA. P*hig*-*higBhigA*-*lacZ* shows little β-gal activity indicating robust HigA repression at P*hig* (∼95% repression). The HigB(L5ext)HigA_2_ variant shows similar repression as wild-type HigB_2_HigA_2_ suggesting there is no difference between a trimeric or tetrameric HigBHigA complex.

**Figure 8.**
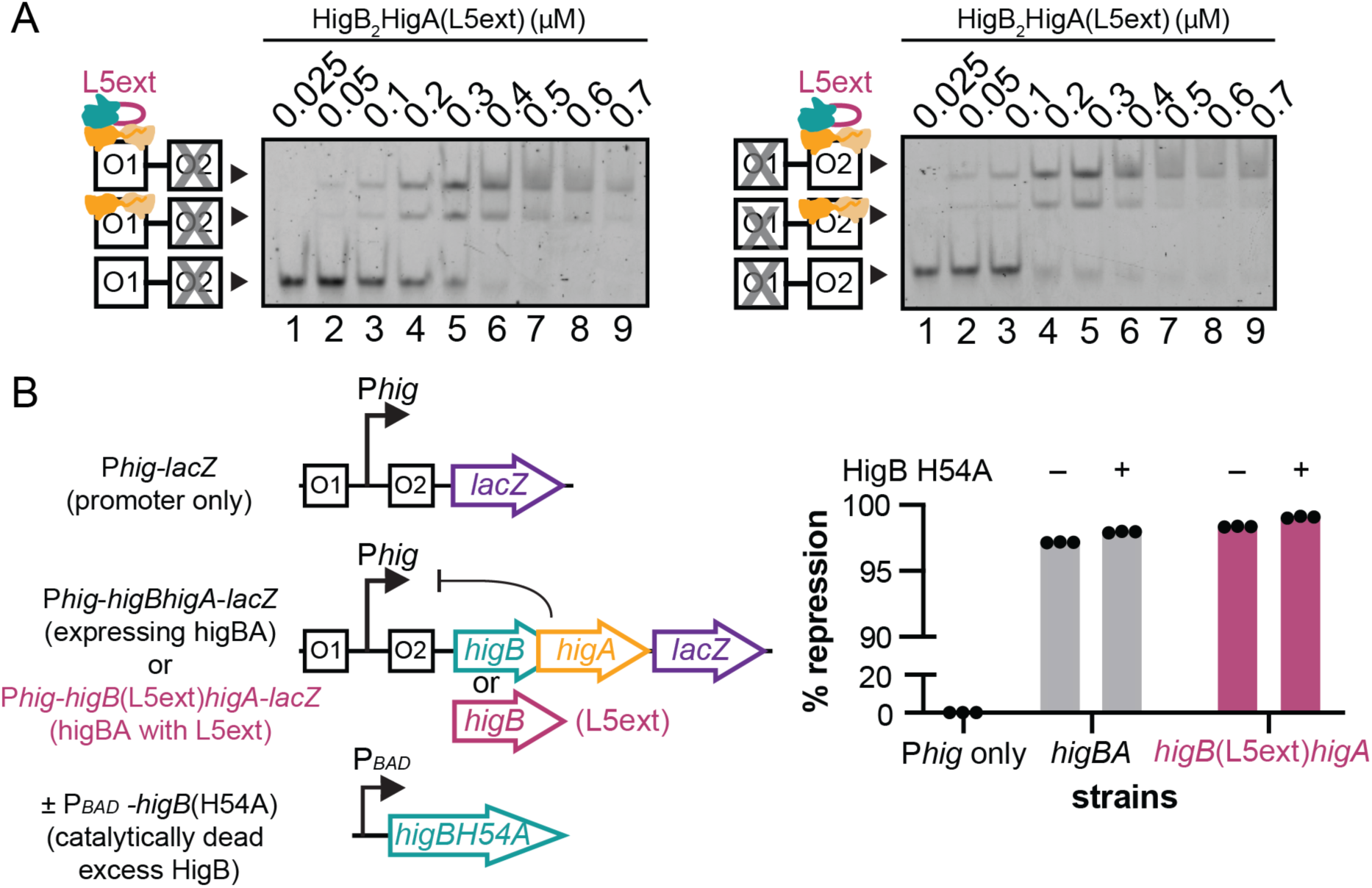
Trimeric HigB(L5ext)HigA_2_ is sufficient to repress transcription of P*hig*. (**A**) EMSA of HigB(L5ext)HigA_2_ binding to O1 DNA (O2 scrambled; *left*) or to O2 DNA (O1 scrambled; *right*). (**B**) β-gal assays of *E. coli* BW25113 transformed with either pQF50-P*hig*-*lacZ* (P*hig* only), pQF50-P*hig*-*higBhigA*-*lacZ* (*higBhigA*), pQF50-P*hig*-*higB*(L5ext)*higA*-*lacZ* (*higB*(L5ext)*higA*), and/or pBAD33-*higB*(H54A). P*hig* only demonstrates the maximum amount of β-gal activity (black bar, 0% repression). Constructs containing either a wild-type HigBHigA (grey bars) or a HigB(L5ext)HigA variant (pink bars) both repress transcription (first bar in each group). Excess HigB expression (using a catalytically inactive H54A variant) results in little difference in repression (second bars). Error bars represent the mean ± SD of values of three independent experiments (raw values shown as dots).

One possibility that we wanted to explore was whether *hig* is responsive to changing toxin levels *in vivo* considering that toxin overexpression in the *phddoc*, *ccdBccdA* and *relEB* systems can relieve repression (17, 18, 27). For this assay, we used a HigB variant (H54A) that is not catalytically active so cell growth won’t be impacted by its expression and we know this variant is expressed as detected by Western blot analysis (34, 43). Further, since the active site of HigB is on the opposite surface from its HigA binding surface (26), this variant should not interfere with HigA binding. Overexpression of HigB(H54A) showed minimal impact on repression of P*hig*-*higBhigA* indicating free HigB(H54A) is unable to interact with HigB_2_HigA_2_ bound at O1 or O2 (97.2% vs. 98.0%; **Fig. 8B**). In the case of the engineered HigB(L5ext)HigA_2_, excess HigB(H54A) expression also has no effect on repression and does not interfere with HigA-mediated repression (**Fig. 8B**). These data support a model whereby *hig* repression is only relieved by HigA proteolysis.

## DISCUSSION

The roles of bacterial TA modules have been controversial owing to experimental errors in the construction of *E. coli* TA deletion strains and the ambiguity over what activates toxin expression, antitoxin proteolysis and the release of toxin (10, 44, 45). While these activities are still under debate, the way these modules are transcriptionally autoregulated is known to clearly contribute to their changing expression patterns in response to external stimuli although many outstanding questions still remain (17, 18, 27). One question is how different oligomeric TA complexes influence physical interactions with their DNA operators and the assembly and/ or cooperativity of TA complexes bound at adjacent operator sites. In this study, we focused on the regulation of the *higBhigA* TA module first identified on the antibiotic-resistance Rts1 plasmid associated with a urinary tract infection caused by *P. vulgaris* (32). Our prior work revealed that while the HigB toxin adopts a canonical ribonuclease fold similar to other members of the RelE family (26), HigA antitoxin suppresses HigB activity in an unusual manner, suggesting that transcriptional repression may also be different. We therefore sought to understand how the *higBhigA* operon is regulated and how its diverse architecture might influence its negative regulation of transcription.

Higher-order oligomeric complexes of the PhDDoc, RelBRelE and CcdBCcdA TA complexes alter their thermodynamic interactions with operators and thus influence transcriptional responsiveness (17, 18, 27). We find that although HigA binding at adjacent operators is not cooperative (38) (**Fig. 2**), the addition of HigB promotes the formation of higher-order complexes and disassembly of the complex does appear to occur once a threshold of excess toxin to antitoxin is reached (**Figs. 2,3**). On the surface, these data appear to be consistent with how other TAs are regulated via the conditional cooperativity model or, as other transcriptional systems are described, as molecular rheostats (14). However, in an attempt to perturb the system *in vivo* by increasing HigB concentrations in the presence of the HigBHigA complex bound at O1 and O2 operators, we find no observable change in repression in contrast to these TA systems (**Fig. 8B**). These conflicting data indicate that the perceived disruption of the higher order HigBHigA-O2 DNA complex via EMSA may not be reflective of transcriptional repression that we observe *in vivo*. Further, these results suggest a different mode of regulation for the *higBhigA* operon.

One hallmark of conditional cooperativity that defines transcriptional regulation of TA operons is that the addition of excess toxin to antitoxins bound at operator sites changes the affinity of these interactions because toxin can act both as a co-repressor and de-repressor (17, 18, 22). In general, when the toxin and antitoxin are expressed at equimolar ratios, the toxin functions as a co-repressor and transcriptional repression is enhanced. Once toxin concentrations exceed that of the antitoxin, however, disruption of the repressor complex ensues, antitoxin disengages from the promoter, and transcription proceeds (**Fig. 1**). To achieve this mode of regulation, defined oligomeric states of TA complexes have different affinities: for example, although the structure of RelBRelE complex indicates the complex is tetrameric with two RelE toxins and two RelB antitoxins (21), a trimeric version of the complex (two RelBs, one RelE) is proposed to have the highest affinity for the *rel* operon (18, 21). In the presence of a greater excess of toxin, the tetrameric RelB_2_RelE_2_ forms at adjacent operators causing a change from a high affinity to a low affinity state, and the RelBRelE complex no longer binds to DNA (**Fig. 1**). This model permits the system to be responsive to changing levels of toxin but at this point, there is limited biochemical and structural data that corresponds to these changing oligomeric states.

We serendipitously solved two different structures of the HigBHigA-O2 complexes that differ in their molar ratios of HigA antitoxin to the HigB toxin (**Figs. 4,5**). These different oligomeric states capture, for the first time, how both the tetrameric HigB_2_HigA_2_ and trimeric HigBHigA_2_ interact with O2 to repress transcription. The trimeric HigBHigA_2_-O2 DNA structure was especially unexpected given that the tetrameric HigB_2_HigA_2_ form predominates in the absence of operator (26). The molecular interactions of each HigBHigA complex with its operator are largely maintained, initially suggesting that different oligomeric states may not contribute to changes in repression for this system. Molecular dynamic simulations of both the trimeric and tetrameric HigBHigA-O2 complexes show each complex have similar dynamics when bound to DNA, offering further support for the ability of both oligomeric states to contribute to repression (**Fig. 6**). Further, engineering of a forced trimeric HigBHigA_2_ oligomeric complex revealed similar levels of transcriptional repression (**Figs. 7,8; Fig. S7**). Together, these data support a model where *hig* is regulated as a simple on/off switch incalcitrant to changing levels of toxin and influenced solely by HigA proteolysis likely as result of the Rts1 plasmid not being inherited.

There are several other reasons for why *hig* may be regulated in a different manner from other TA operons. In TA operons regulated as molecular rheostats including *phddoc*, *relBrelE* and *ccdBccdA*, a single promoter controls expression of the operon and the antitoxin is encoded first (17, 18, 22). It has been suggested this operon organization allows for the antitoxin to be expressed in excess of the toxin, which is required to suppress toxicity when the system is transcriptionally de-repressed (46, 47). In *hig*, there is a second weak promoter, P*higA*, that allows for independent expression of HigA to ensure excess HigA is always present to suppress HigB (48). It is not clear if this change in operon organization accounts for why *hig* is regulated via a simple on/off switch but other TA operons such as *mqsRA* that also have this reverse architecture and contain multiple promoters do not appear to be regulated via conditional cooperativity (29). In contrast, the DinJ-YafQ TA module does not have this reverse architecture but does not appear to be regulated by conditional cooperativity (30). Therefore, there must be some other currently unappreciated mechanism by which this TA system balances antitoxin expression for its regulation.

The results presented here provide new insight into the transcriptional regulation of the plasmid-associated *hig* operon and add to the growing diversity of mechanisms used to balance transcriptional responses of these abundant bacterial gene pairs. In the future, additional biophysical studies are needed to reconcile the role of changing macromolecular complex formation in the regulation of TA pairs and to align these properties with transcriptional responsiveness.

## MATERIALS AND METHODS

### Strains and plasmids

*E. coli* BL21(DE3) cells were used for expression of His_6_-HigA, His_6_- HigBHigA and HigBHigA-His_6_ proteins from pET28a, pET28a and pET21c vectors, respectively as previously reported (26). *E. coli* BW25113 cells were used for all β-gal experiments and HigB(H54A)-His_6_ expression (49). All point mutations were introduced by site-directed mutagenesis and sequences were verified by DNA sequencing (Genewiz).

### HigA, HigB and HigBHigA expression and purification

The His_6_-HigA, His_6_-HigBHigA and HigBHigA-His_6_ protein complexes were overexpressed and purified as previously described with minor modifications (26). These differences included incubation of His_6_-HigA at 18°C overnight after protein induction and removal of the His_6_ tag from His_6_-HigA and His_6_- HigBHigA with thrombin prior to gel filtration chromatography. HigB(H54A) protein was overexpressed and purified as previously described (43).

### Electrophoretic mobility shift assays (EMSAs)

To construct the dsDNA for the EMSA, pairs of complementary single-stranded oligonucleotides were diluted to 2 μM each in 100 mM NaCl, 10 mM Tris-HCl pH8. The O1-O2, O1-O2(scrambled) or O1(scrambled)-O2 oligonucleotide mixtures (**Table S1**) of the *hig* promoter fragment were incubated in boiling water and then cooled at room temperature overnight. The dsDNA oligos were diluted to 150 nM in EMSA binding buffer (100 mM NaCl, 10 mM MgCl_2_, 5% glycerol, 0.01 mg/mL bovine serum albumin). Purified wild-type HigA, HigBHigA, and HigB(L5ext)HigA proteins were diluted to 10 μM in EMSA binding buffer and serially diluted to give a series of protein concentrations ranging from 25 nM to 0.8 μM. The binding reactions were incubated on ice for 20 min and 10 μL of each reaction was loaded onto 8% native, polyacrylamide-0.5X TBE/10% glycerol gels (50 mM Tris-HCl pH 8, 50 mM boric acid, 5 mM EDTA, 10% glycerol) and subjected to electrophoresis at 110 V limiting on ice for 60 min. To visualize the DNA and DNA-protein complexes, the gels were stained with SYBR green nucleic acid gel stain (ThermoFisher Scientific) in 0.5X TBE/10% glycerol for 30 min with gentle agitation and then the fluorescence was imaged with a Typhoon Trio phosphoimager (GE Healthcare; 488 nm excitation and 526 nm emission). Assays were performed in duplicate with representative gels shown. Band intensities for both free and bound *hig* DNA were quantified with ImageQuant 1D gel analysis software using the rolling ball background subtraction. For HigA or HigBHigA bound to either O1 or O2, the binding data were fit using a one site-specific binding equation (Y (specific binding, μM) = Bmax * X / [K_D_ + X]) in GraphPad Prism 9.0.0.

### Crystallization, data collection and structure determination of the HigBHigA-O2 DNA complexes

The complex was formed by mixing either His_6_-HigBHigA or selenomethionine-derivatized HigBHigA-His_6_ (both in 40 mM Tris-HCl, pH 7.5, 250 mM KCl, 5 mM MgCl_2_, and 5 mM β-mercaptoethanol) with O2 operator DNA (10 mM Tris, pH 8, 100 mM NaCl, and 1 mM EDTA) at one HigB_2_HigA_2_ tetramer to one O2 dsDNA molar ratio. The complexes were diluted to 5.95 mg/mL HigBHigA and 1.55 mg/mL O2 DNA by the addition of buffer (20 mM Tris, pH 8, 10 mM MgCl_2_, and 100 mM NaCl). Crystals of HigBHigA bound to O2 DNA were grown by sitting drop vapor diffusion and crystallized in 0.2 M CaCl_2_ and 10-25% (w/v) polyethylene glycol 3,350 at 20°C. Both crystal forms grew after two days and were cryoprotected by serially increasing the concentration of ethylene glycol in the mother liquor from 10-30% (w/v) followed by flash freezing in liquid nitrogen.

Two X-ray datasets were collected at the Northeastern Collaborative Access Team (NE-CAT) 24-ID-C and Southeast Regional Collaborative Access Team (SER-CAT) 22-ID facilities at the Advanced Photon Source (APS) at the Argonne National Laboratory. For the tetrameric HigB_2_HigA_2_-O2 complex, 360° of data (0.5° oscillations) were collected on a PILATUS 6M-F detector (DECTRIS Ltd., Switzerland) using 0.9792 Å radiation. For the trimeric HigBHigA_2_-O2 complex, 90° of data (0.5° oscillations) were collected on a MARMOSAIC 300 mm CCD detector (Rayonix, L.L.C., USA) using 1.0 Å radiation. XDS was used to integrate and scale the data (50). The tetrameric HigB_2_HigA_2_-O2 structure was solved by single wavelength anomalous diffraction phasing using AutoSol from the PHENIX software suite (51) and thirteen heavy atom sites were found. The trimeric HigBHigA_2_-O2 structure was solved using the structure of the HigA_2_ dimer (PDB code 6CF1) as a molecular replacement search model in the PHENIX software suite. XYZ coordinates, real space, and B-factors (isotropic) were refined iteratively in PHENIX and model building was performed using the program Coot (52). Final refinement of the structures gave crystallographic R_work_/R_free_ of 17.6/21.8% for trimeric HigBHigA_2_-O2 and 17.5/22.1% for tetrameric HigB_2_HigA_2_-O2. All figures were created in PyMol (53).

### β-galactosidase assays

The *hig* operon was chemically synthesized (IDT), digested and ligated into a pQF50 vector with *lacZ* downstream (pQF50-*hig* constructs). *E. coli* BW25113 transformed with pQF50-*hig* variants or pBAD33-*higB*(H54A) were used for all experiments. Two methods were used to perform β-gal assays. The first set of β-gal assays presented herein (with constructs containing wild-type versus mutated operators) were performed using the PopCulture® Reagent based method (54). All overnight cultures were grown in M9 minimal media supplemented with 0.2% glucose, 1 M MgSO_4_, 1 M CaCl_2_, and 10% casamino acids. Subsequent experiments were performed with M9 minimal media supplemented with 20% glycerol instead of glucose. OD_600_ was measured hourly until an OD_600_ of 0.2 was reached, arabinose was added to a final concentration of 0.2%, and further incubated for 4 hrs. 1 mL aliquots were pelleted, resuspended in 500 µL of M9 media and diluted to an OD_600_ of 0.5 in M9 media. 80 µL were transferred to 96-well plate and 120 µL of freshly mixed β-gal reagent (60 mM Na_2_SO_4_, 40 mM NaH_2_PO_4_, 10 mM KCl, 1 mM MgSO_4_, 36 mM β-me, 166 µL/mL T7 lysozyme, 1.1 mg/mL ortho-Nitrophenyl-β-galactoside and 6.7% PopCulture® Reagent (Millipore-Sigma)) was added to each well. 80 µL of M9 media was also added to wells containing 120 μL of β-gal reagent as a negative control. The microplate was read by a Biotek Cytation 5 multi-mode reader pre-incubated at 30°C, with OD_600_ and OD_420_ measurements taken every 5 min for 1 hr with agitation.

The second set of β-gal assays presented herein (in the absence or presence of HigB(H54A)) were performed using a method previously described (38). In both approaches, activity in Miller Units (M.U.) was measured using the formula: total activity (M.U.) = (1000*OD_420_) / (OD_600_ * volume of culture used (mL) * 0.5). Assays were performed in triplicate with two technical replicates.

### Differential scanning fluorimetry (DSF)

The thermal stability of wild-type HigBHigA and HigB(L5ext)HigA were assessed using a Tycho NT.6 instrument (NanoTemper). Protein was heated at 0.1°C steps over a temperature range of 35°C to 95°C, during which intrinsic fluorescence at 350 and 330 nm was measured. Inflection temperature (T_i_) was determined for each apparent unfolding transition from the temperature-dependent change in the ratio of 350 and 330 nm measurements. Assays were performed in triplicates.

### Molecular dynamics simulations

Starting models for molecular dynamics (MD) simulations were prepared from PDB codes 6W6U (HigB_2_HigA_2_-O2) and 6WFP (HigBHigA_2_- O2). Simulations were performed on the tetrameric or trimeric HigBHigA structures in the absence or presence of O2 DNA (HigB_2_HigA_2_-O2, HigBHigA_2_-O2, HigB_2_HigA_2_ and HigBHigA_2_). All complexes were prepared using the Xleap module of AmberTools 18 (55), the ff14SB forcefield for protein atoms (56) and the OL15 forcefield (57) for DNA. Complexes were solvated in an octahedral box of TIP3P water (58) with a 10 Å buffer. Ions were added to each complex to achieve a final concentration of 150 mM NaCl. Minimization was performed in three rounds, each employing steepest descent (5000 steps) followed by conjugate gradient (5000 steps). In the first round, restraints of 500 kcal/mol-Å^2^ were applied to all solute atoms. In the second round, solute restraints were reduced to 100 kcal/mol-Å^2^. All restraints were removed in the third round. Complexes were heated from 0 to 300 K with a 100-ps run with constant volume periodic boundaries and restraints of 10-kcal/mol-Å^2^ on solute atoms. All MD simulations were performed using AMBER2018 (55, 59, 60). Two stages of equilibration were performed: 10 ns MD in the NPT ensemble with 10-kcal/mol-Å^2^ restraints on solute atoms, followed by an additional 10 ns MD run with restraints reduced to 1 kcal/mol-Å^2^. Finally all restraints were removed and 1 microsecond production simulations obtained for each complex. Long-range electrostatics were evaluated with a cutoff of 10 Å and all heavy atom-hydrogen bonds were fixed with the SHAKE algorithm (61). Following MD, the CPPTRAJ module (62) of AmberTools 18 was used to calculate root mean square fluctuations (RMSF) of each protein reside in each complex.

## ACKNOWLEDGEMENTS

Research reported in this publication was partially supported by a National Science Foundation CAREER award MCB 0953714 (CMD), a NIH Biochemistry, Cellular and Molecular Biology Graduate Training Grant 5T32GM8367 (MAS), and NIH NRSA F31 Fellowship GM108351 (MAS) and a Burroughs Wellcome Fund Investigator in the Pathogenesis of Infectious Disease award (CMD). We thank F. M. Murphy IV and staff members of the NE-CAT beamlines for assistance during data collection and G. L. Conn and other Dunham lab members for critical reading of the manuscript. This work is based upon research conducted at the Northeastern Collaborative Access Team beamlines, which are funded by the National Institute of General Medical Sciences from the National Institutes of Health (P30 GM124165). Additional data were collected at the Southeast Regional Collaborative Access Team (SER-CAT) 22-ID (or 22-BM) beamline at the Advanced Photon Source, Argonne National Laboratory. SER-CAT is supported by its member institutions (www.ser-cat.org/members.html) and equipment grants (S10RR25528 and S10RR028976) from the National Institutes of Health. Use of the Advanced Photon Source was supported by the U. S. Department of Energy, Office of Science, Office of Basic Energy Sciences, under Contract No. DE-AC02-06CH11357.

